# Numerical and analytical simulation of the growth of amyloid-β plaques

**DOI:** 10.1101/2023.09.11.557187

**Authors:** Andrey V. Kuznetsov

**Affiliations:** Department of Mechanical and Aerospace Engineering, North Carolina State University, Raleigh, NC 27695-7910, USA

**Keywords:** neuron, Alzheimer’s disease, Finke−Watzky model, mathematical modeling, polymerization, cube root hypothesis

## Abstract

Numerical and analytical solutions were employed to calculate the radius of an amyloid-β (Aβ) plaque over time. To the author’s knowledge, this study presents the first model simulating the growth of Aβ plaques. Findings indicate that the plaque can attain a diameter of 50 μm after 20 years of growth, provided the Aβ monomer degradation machinery is malfunctioning. A mathematical model incorporates nucleation and autocatalytic growth processes using the Finke-Watzky model. The resulting system of ordinary differential equations was solved numerically, and for the simplified case of infinitely long Aβ monomer half-life, an analytical solution was found. Assuming that Aβ aggregates stick together and using the distance between the plaques as an input parameter of the model, it was possible to calculate the plaque radius from the concentration of Aβ aggregates. This led to the “cube root hypothesis,” positing that Aβ plaque size increases proportionally to the cube root of time. This hypothesis helps explain why larger plaques grow more slowly. Furthermore, the obtained results suggest that the plaque size is independent of the kinetic constants governing Aβ plaque agglomeration, indicating that the kinetics of Aβ plaque agglomeration is not a limiting factor for plaque growth. Instead, the plaque growth rate is limited by the rates of Aβ monomer production and degradation.

## 1. Introduction

Alzheimer’s disease (AD) is a devastating neurodegenerative disorder, impacting nearly 50 million individuals globally. While recent FDA-approved therapies may decelerate its progression, there are currently no definitive cures for AD [1-4]. It is crucial to develop mechanistic models of processes involved in AD to pave the way for potential future treatments.

Senile plaques, primarily composed of accumulated amyloid-β (Aβ) peptides, represent a key characteristic of AD [5]. A significant surge of interest in Aβ aggregation arose from the amyloid cascade hypothesis, which posited that the formation of Aβ aggregates initiates all other pathological processes in AD [6]. While this hypothesis has undergone extensive investigation, it has also faced recent criticism from some authors [7-9]. The reason is the failure of many Aβ-targeted clinical trials for AD [10]. Yet, lifelong Aβ reduction may prevent or reduce the risk of dementia [11]. Multi-target drugs, addressing Aβ among other factors, may be promising for AD treatment. Additionally, Aβ serves as a crucial biomarker in AD diagnosis [12-15].

Previous mechanistic models of AD [16-18] primarily focused on elucidating intraneuronal mechanisms contributing to the development of AD, such as the formation of neurofibrillary tangles composed of aggregated tau proteins and the generation of Aβ monomers at the cell membrane. However, these investigations did not simulate the gradual extracellular buildup of Aβ plaques. In contrast, the present study aims to replicate the gradual growth of an Aβ plaque, a process that can span decades in humans. This process involves the generation of amyloid precursor protein (APP) within neurons and its subsequent cleavage by β-and γ-secretases at lipid membranes [14,19-21]. The majority of these newly formed Aβ monomers are released into the extracellular environment, where they hold the potential to aggregate [1]. The goal of this paper is to develop a mechanistic model predicting how the radius of an Aβ plaque increases in time.

## 2. Materials and models

### 2.1 Model equations

The F-W model simplifies the intricate process of protein aggregation into two key steps: nucleation and autocatalysis. During nucleation, new aggregates are continuously generated, while in the autocatalysis step, these aggregates undergo rapid surface growth [22,23]. These two pseudo-elementary reaction steps can be summarized as follows:

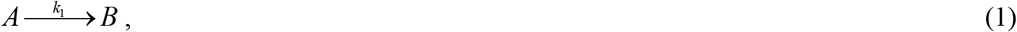

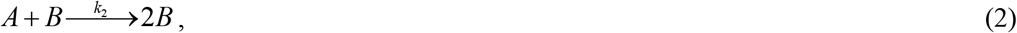

where *A* refers to a monomeric protein, and *B* represents a protein that has undergone amyloid conversion. The kinetic constants, denoted as *k*_1_ and *k*_2_, correspond to the rates of nucleation and autocatalytic growth, respectively [22]. The primary nucleation process, as defined by Eq. (1), exclusively involves monomers. Conversely, secondary nucleation, described by Eq. (2), involves both monomers and preexisting aggregates of the same peptide [24].

In [22], the Finke-Watzky (F-W) model was utilized to fit previously published data on Aβ aggregation [25,26]. Typically, this data is presented for two isoforms, Aβ40 and Aβ42, with the latter being more prone to aggregation. In this context, the F-W model is applied to simulate the conversion of monomers, whose concentration is denoted as *C*_*A*_, into aggregates, whose concentration is denoted as *C*_*B*_ . These aggregates encompass various forms of Aβ oligomers, protofibrils, and fibrils [5]. Aβ aggregates formed through this process assemble in amyloid plaques. It is worth noting that the simplified nature of the F-W model does not allow for differentiation between various aggregate types and sizes. The current model does not simulate the process by which adhesive Aβ fibrils assemble into an Aβ plaque. It is assumed that this assembly occurs faster than the formation of Aβ fibrils.

If soluble Aβ were to diffuse at a slower rate than the pace of aggregation, it would lead to growth primarily occurring at the tips of aggregates, resulting in a branching, tree-like structure. However, this contradicts what has been observed in experiments [27]. The rapid diffusion of Aβ monomers suggests minimal variation in their concentration, *C*_*A*_, between neurons. With this assumption, the control volume (CV) displayed in Fig. 1 with a volume *L*^3^ can be treated as a lumped capacitance body. Expressing the conservation of Aβ monomers within this CV yields the following equation:

**Fig. 1.**
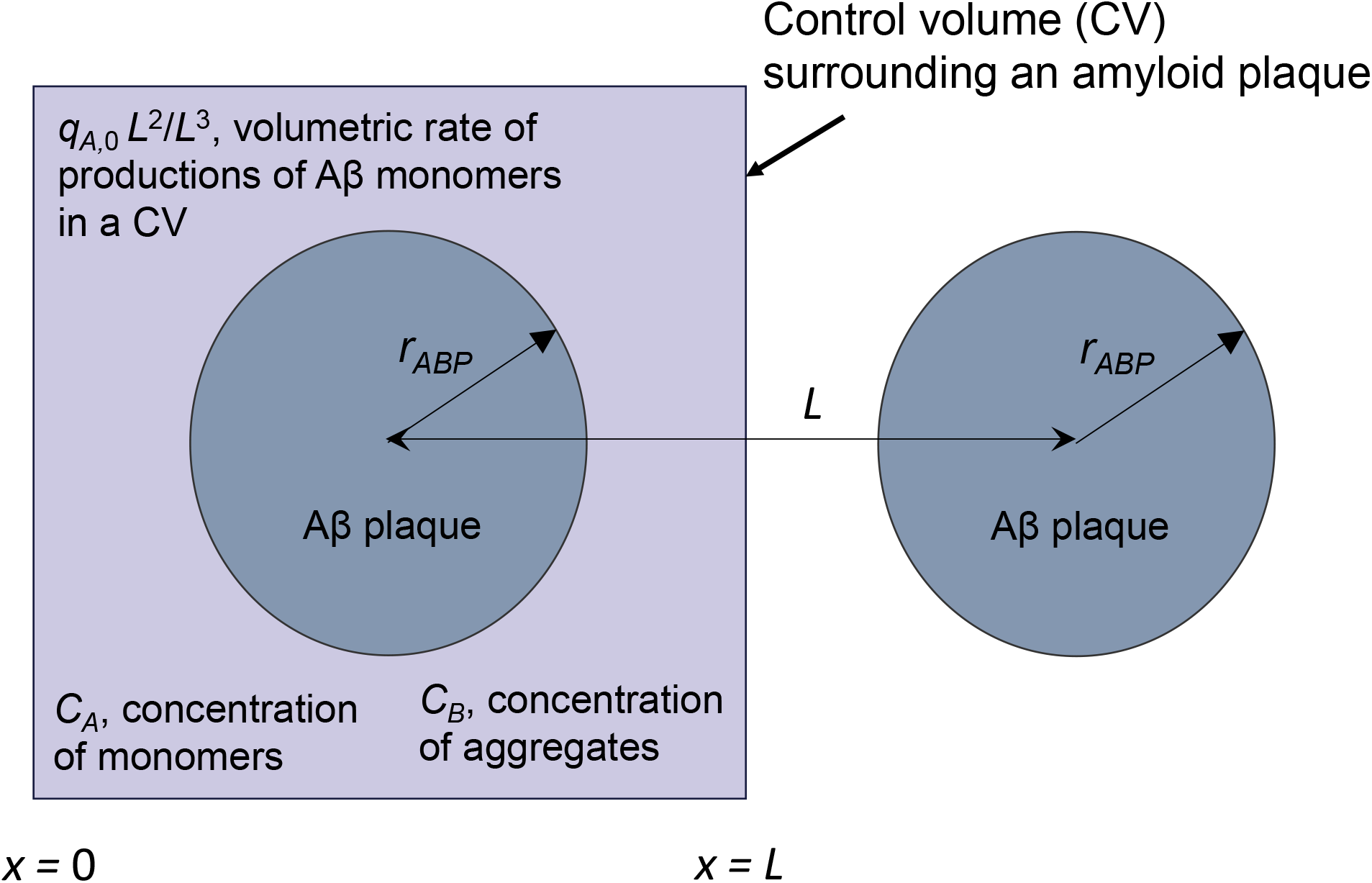
A cubic CV with a side length of *L*containing a single growing Aβ plaque.

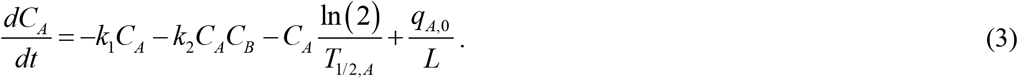

The first term on the right-hand side of Eq. (3) simulates the rate of conversion of Aβ monomers into aggregates, while the second term simulates the rate of this conversion through autocatalytic growth. These expressions for the first and second terms arise from applying the law of mass action to the reactions described by Eqs. (1) and (2), respectively. The third term on the right-hand side of Eq. (3) accounts for the rate at which Aβ monomers degrade. The fourth term represents the rate of Aβ monomer production, as Aβ monomers are continuously generated through the cleavage of APP [28,29]. The production of Aβ monomers is a surface phenomenon that takes place at lipid membranes [5]. Since the precise area of the lipid membrane contributing to the production of Aβ monomers per CV is unknown, the production was normalized by the area of one of the CV’s faces, *L*^2^ . *q*_*A*,0_ should be understood as the average flux of Aβ monomers into the CV per unit area of one of the CV’s faces. The total rate at which monomers enter the CV is *q*_*A*,0_ *L*^2^ . Since the lumped capacitance method is used, the rate of Aβ monomers entering the CV is converted into volumetric monomer generation, providing the same number of monomers per unit time, *q*_*A*,0_ *L*^2^ / *L*^3^ . This is the reason why the fourth term on the left-hand side of Eq. (3) is inversely related to *L*.

If the conservation of Aβ aggregates is written for the CV, the following equation is obtained:

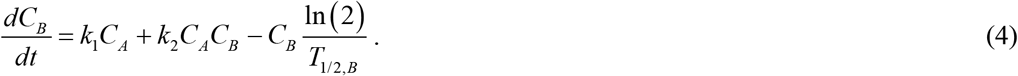

The first term on the right-hand side of Eq. (4) represents the rate at which Aβ aggregates are produced through nucleation, while the second term simulates the rate of their production via autocatalytic growth. These terms have equal magnitudes, but opposite signs when compared to the first and second terms in Eq. (3). This is because, in the F-W model, the rate of aggregate production equals the rate of monomer disappearance. The third term in Eq. (4) represents the rate of Aβ aggregate degradation, with the assumption that this degradation is significantly slower than that of monomers [30].

The initial conditions are as follows:

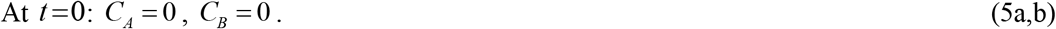

The sole independent variable in the model is time, *t*. The dependent variables are listed in Table 1, while the parameters employed in the model are listed in Table 2.

**Table 1.** Dependent variables utilized in the model.

| Symbol | Definition | Units |
| --- | --- | --- |
| $C_A$ | Concentration of Aβ monomers | $\mu\text{M}$ |
| $C_B$ | Concentration of Aβ aggregates | $\mu\text{M}$ |
| $r_{ABP}$ | Radius of a growing Aβ plaque | $\mu\text{m}$ |

**Table 2.**
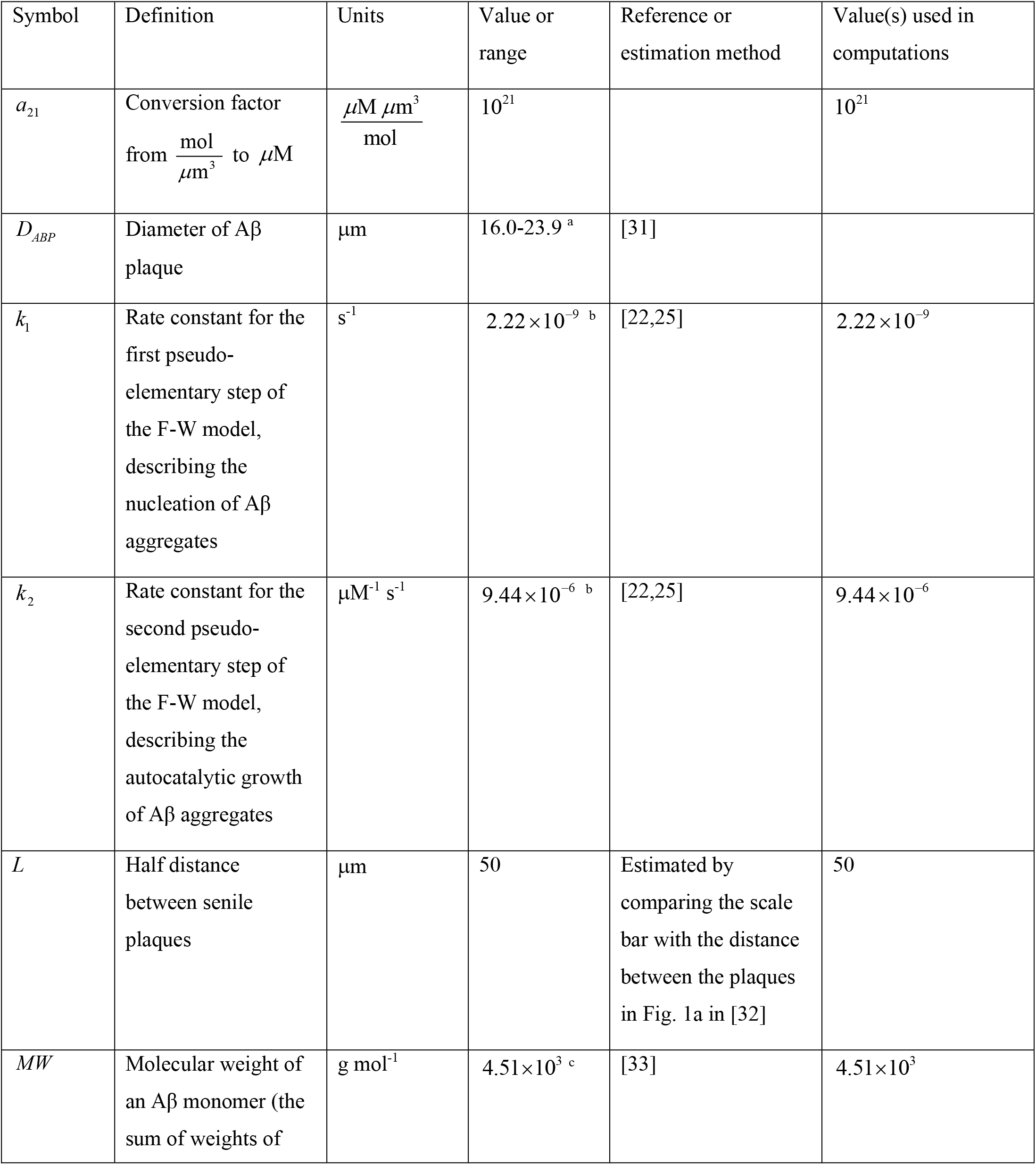
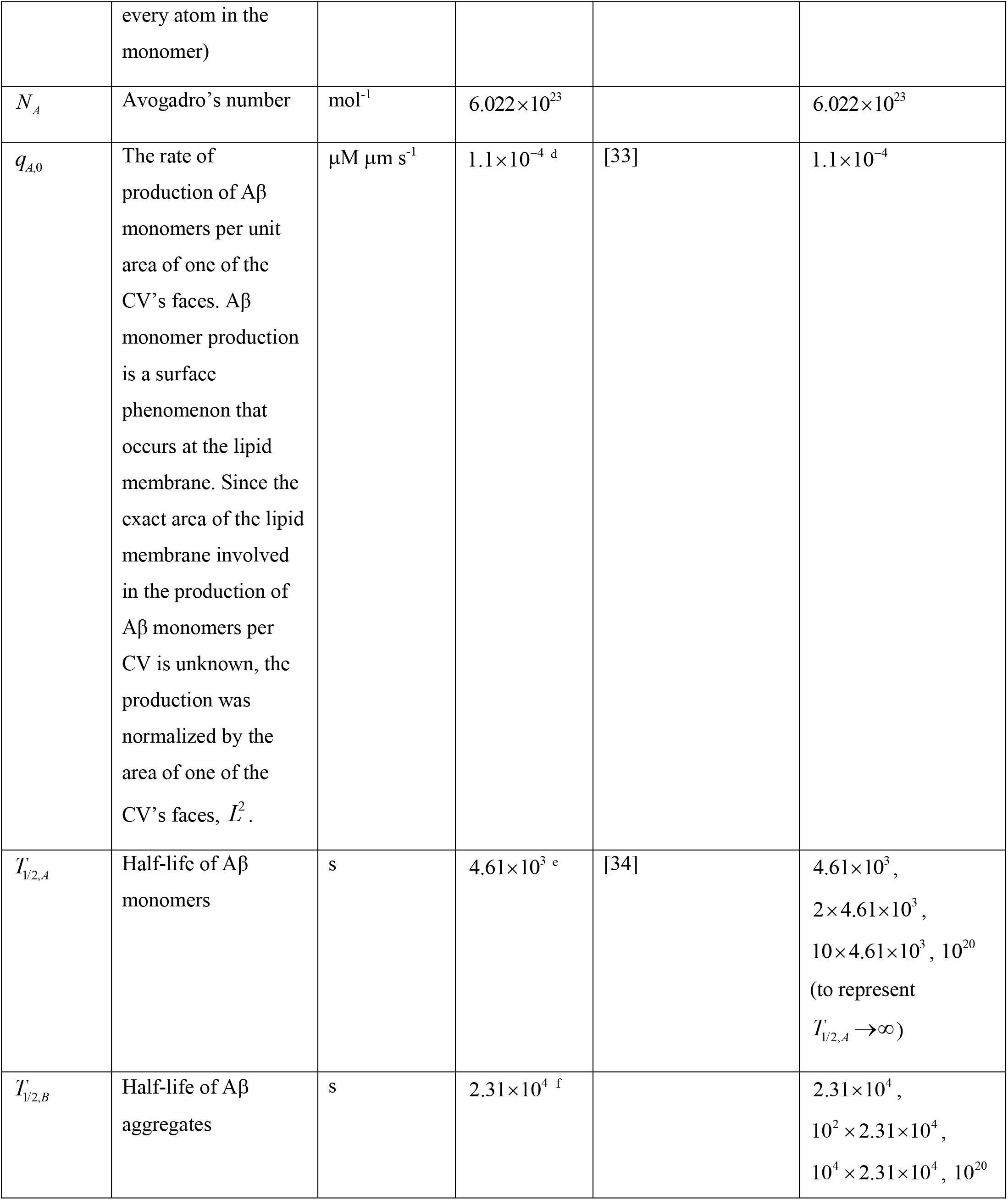
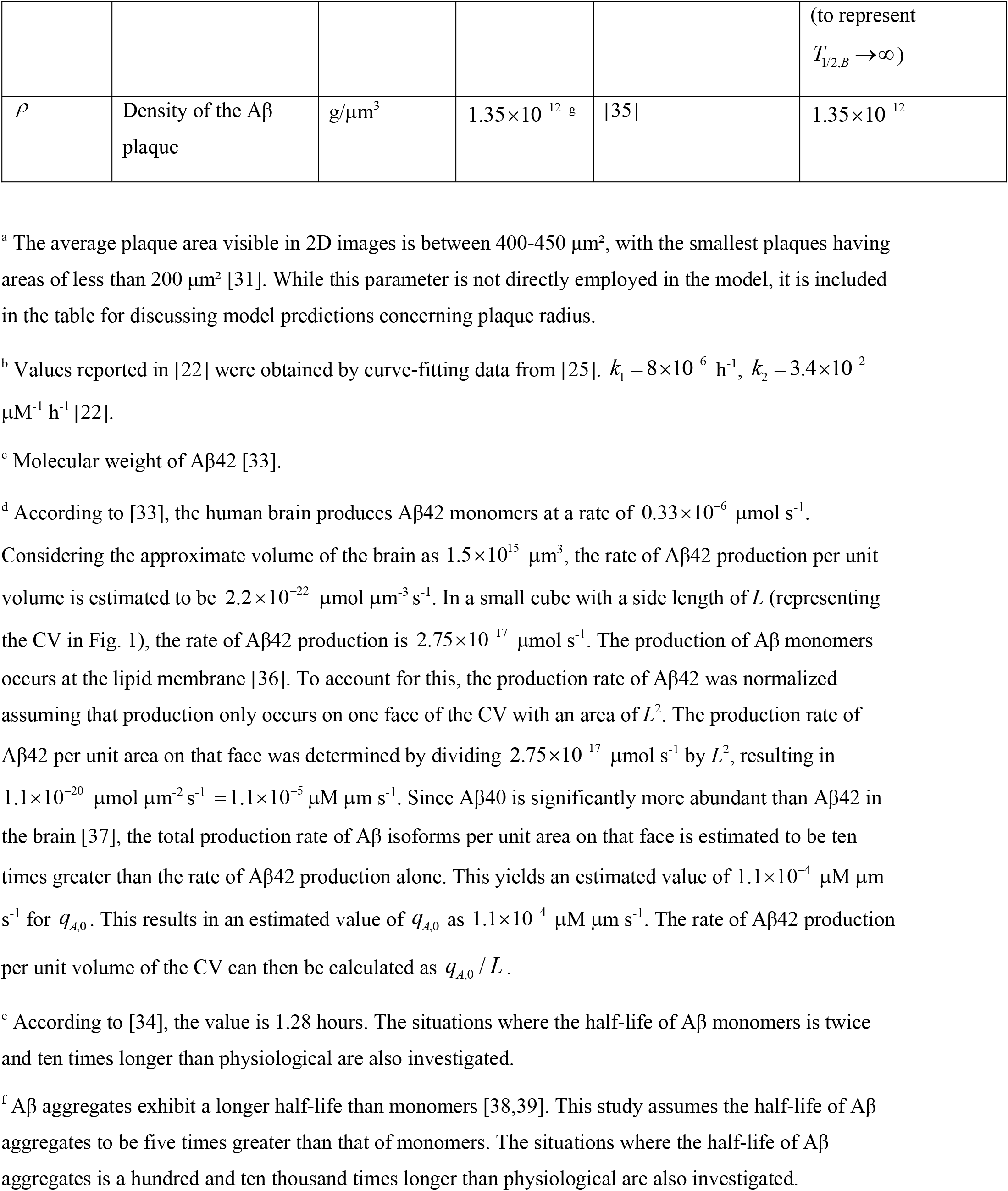
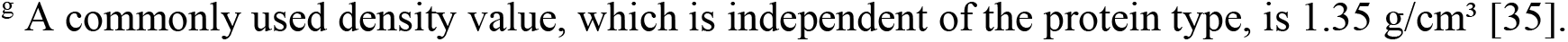
Model parameters and their estimated values.

The dimensionless form of Eqs. (3) and (4) can be expressed as follows:

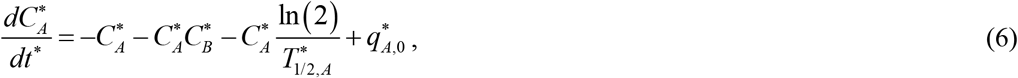

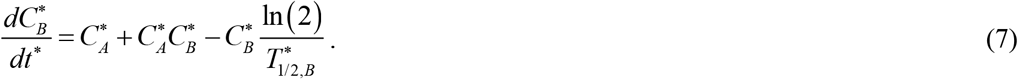

The dimensionless form of the initial conditions defined by Eq. (5) is as follows:

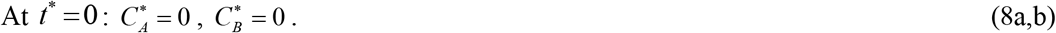

Table 3 presents the dimensionless independent variable, while Table 4 summarizes the dimensionless dependent variables. Table 5 provides the dimensionless parameters used in the model.

**Table 3.** Dimensionless independent variable in the model.

| Symbol | Definition |
| --- | --- |
| $t^*$ | $t k_1$ |

**Table 4.** Dimensionless dependent variables employed in the model.

| Symbol | Definition |
| --- | --- |
| $C_A^*$ | $C_A k_2 / k_1$ |
| $C_B^*$ | $C_B k_2 / k_1$ |
| $r_{ABP}^*$ | $r_{ABP} / L$ |

**Table 5.** Dimensionless parameters utilized in the model.

| Symbol | Definition | Value(s) obtained using the dimensional parameter values listed in the last column of Table 2 |
| --- | --- | --- |
| $q_{A,0}^*$ | $q_{A,0}k_2 / (k_1^2L)$ | $4.21 \times 10^6$ |
| $T_{1/2,A}^*$ | $T_{1/2,A}k_1$ | $1.02 \times 10^{-5}$, $2 \times 1.02 \times 10^{-5}$, $10 \times 1.02 \times 10^{-5}$, $2.22 \times 10^{11}$ (to represent $T_{1/2,A}^* \rightarrow \infty$) |
| $T_{1/2,B}^*$ | $T_{1/2,B}k_1$ | $5.12 \times 10^{-5}$, $10^2 \times 5.12 \times 10^{-5}$, $10^4 \times 5.12 \times 10^{-5}$, $2.22 \times 10^{11}$ (to represent $T_{1/2,B}^* \rightarrow \infty$) |
| $\rho^*$ | $\frac{\rho}{MW} a_{21} \frac{k_2}{k_1}$ | $1.27 \times 10^9$ |

Eqs. (6) and (7) constitute a pair of ordinary differential equations. A well-validated MATLAB’s ODE45 solver (MATLAB R2020b, MathWorks, Natick, MA, USA) was employed to solve these equations subject to initial conditions given by Eq. (8). To ensure accuracy of the solution, the error tolerance parameters, RelTol and AbsTol, were set to 1e-10.

When Eqs. (6) and (7) were added together, the following result was obtained:

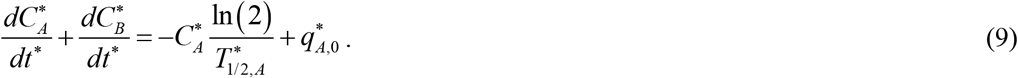

By integrating Eq. (9) with respect to time, assuming *T*_1/2,*A*_ ⟶∞ and *T*_1/2,*B*_ ⟶∞, and utilizing the initial condition given by Eq. (8), the following result was obtained:

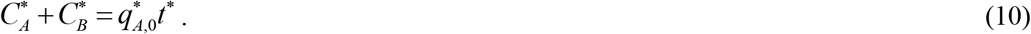

The increase in 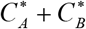 over time is attributed to the continuous generation of Aβ monomers.

By eliminating 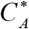 from Eq. (7) using Eq. (10), the following was obtained:

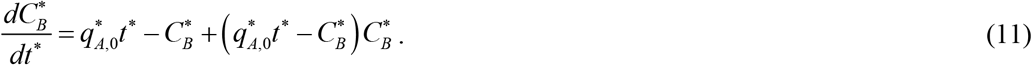

Eq. (11) is similar to the one examined in [40]. To find the exact solution of Eq. (11) with the initial condition given by Eq. (8b), the DSolve function followed by the FullSimplify function in Mathematica 13.3 (Wolfram Research, Champaign, IL) were utilized. The resulting solution is presented below:

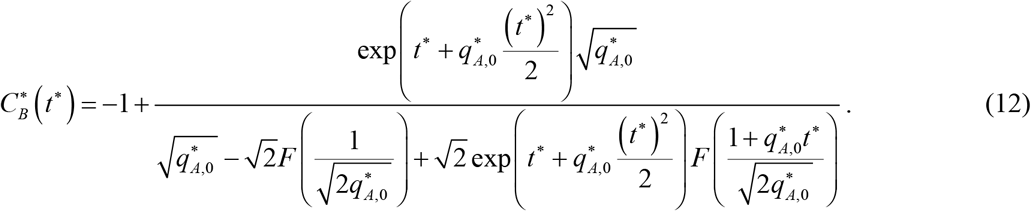

In this equation, *F* ( *x* ) represents Dawson’s integral:

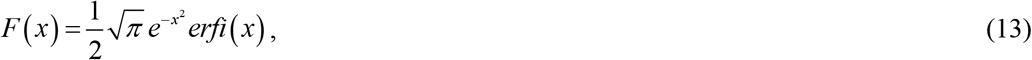

where *erfi* ( *x*) is the imaginary error function.

If *t*^*^ ⟶ 0, Eq. (12) implies that

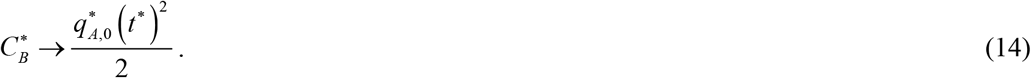

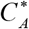can then be determined using Eq. (10) as:

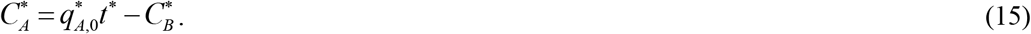

The exact solution given by Eqs. (12) and (15) is quite complex. A more elegant approximate solution, which is also valid when *T*_1/2,*A*_ ⟶∞ and *T*_1/2,*B*_ ⟶∞, is as follows:

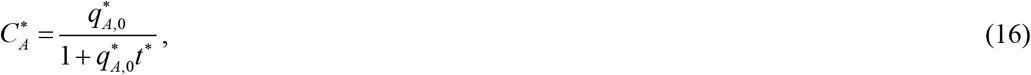

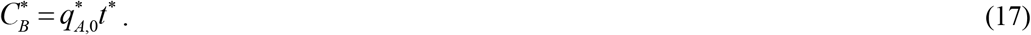

As *t*^*^ ⟶∞, Eq. (16) predicts that 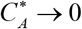. It is important to note that Eqs. (16) and (17) are useful for larger times. However, they are not applicable when *t*^*^ ⟶ 0 ; for instance, Eq. (16) yields 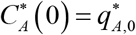, which contradicts the initial condition specified in Eq. (8a).

The Aβ plaque’s growth (Fig. 1) is simulated using the following approach. The total number of Aβ monomers that are integrated into the Aβ plaque, denoted as *N*, at time *t*, is determined using the following equation [41]:

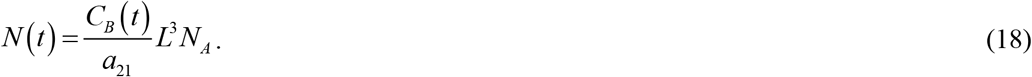

Alternatively, *N* (*t* ) can be calculated as described in [41]:

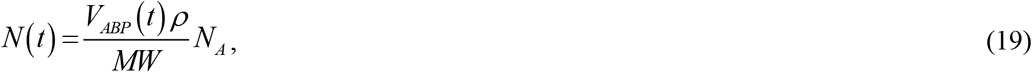

where *MW* denotes the average molecular weight of an Aβ monomer.

By equating the expressions on the right-hand sides of Eqs. (18) and (19) and solving for the volume of an Aβ plaque, the following result is obtained:

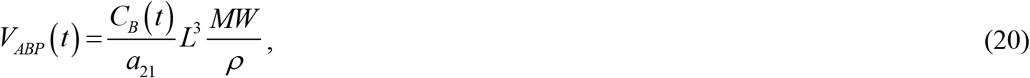

where 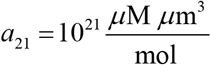 denotes the conversion factor from 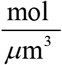 to *μ*M .

Assuming that the Aβ plaque is spherical in shape,

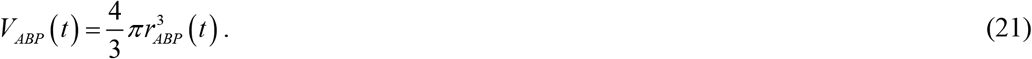

Setting the right-hand sides of Eqs. (20) and (21) equal and solving for the radius of the Aβ plaque, the following is obtained:

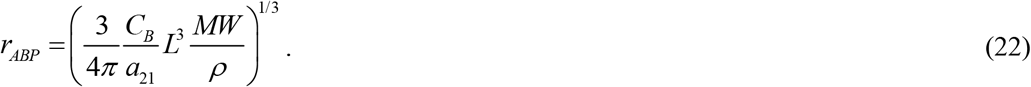

Expressed in dimensionless variables, Eq. (22) becomes:

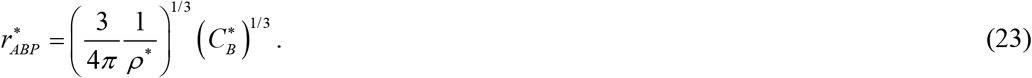

Substituting Eq. (17) into Eq. (23) yields the following approximate solution for the dimensionless radius of the Aβ plaque:

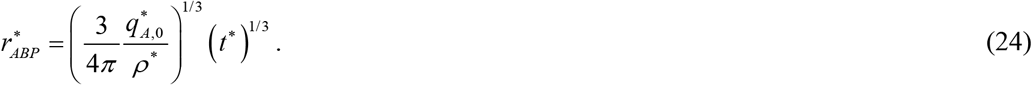

Returning to the dimensional variables, the following equation is obtained:

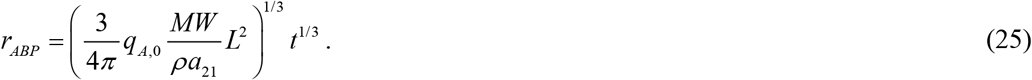

According to Eq. (25), the size of Aβ plaques increases proportionally to the cube root of time. One can infer from Eq. (25) that the rate of Aβ plaque growth is

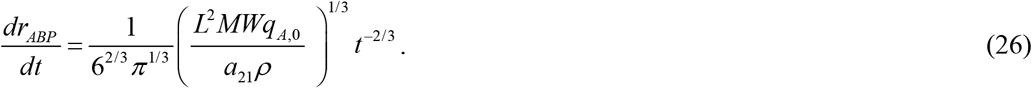

Eq. (26) offers an explanation for the deceleration of Aβ plaque growth over time. This implies that smaller (younger) plaques tend to undergo faster growth compared to larger plaques, a phenomenon observed in previous research [42].

If *t*^*^ ⟶ 0, substituting Eq. (14) into Eq. (23) yields the following approximate solution for the dimensionless radius of the Aβ plaque:

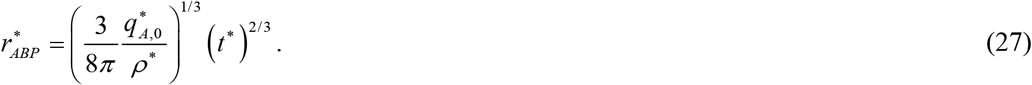

Returning to the dimensional variables, the following equation is obtained:

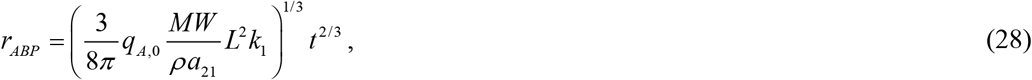

which suggests that at small times, the radius of an Aβ plaque depends on the kinetic constant that describes the nucleation of Aβ aggregates.

Interestingly, Eq. (25) predicts that the radius of an Aβ plaque is independent of the kinetic constants describing the rate of Aβ plaque agglomeration, *k*_1_ and *k*_2_ . However, Eq. (25) is valid only at large times.

At small times, the plaque growth is described by Eq. (28), which suggests that the initiation of the plaque formation process is driven by kinetic factors. The kinetic-driven initial stage of plaque growth agrees with the fast appearance of plaques in a mouse model observed in [43]. However, the continued growth of the plaques is restricted by the rate of Aβ monomer production and, if the half-lives of the monomers and aggregates are finite, by the rate at which the monomers and aggregates degrade. The finding that plaque growth is restricted by Aβ peptide production at extended times is consistent with observations reported in [44].

### 2.2 Sensitivity analysis

The sensitivity of a growing Aβ plaque radius, *r*_*ABP*_, to the two parameters in Eq. (25), *L* and *q*_*A*,0_, as well as to *T*_1/ 2, *A*_ was investigated. To do this, the local sensitivity coefficients [45-48] were computed. The sensitivity coefficients represent the first-order partial derivatives of the observable (in this case, the radius of the growing Aβ plaque) to *L, q*_*A*,0_, and *T*_1/ 2, *A*_, for example, 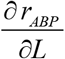. The dimensionless relative sensitivity coefficients [46,49] were then determined using the following equation, for example:

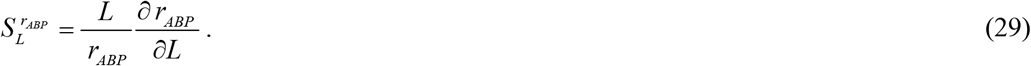

Utilizing Eq. (25):

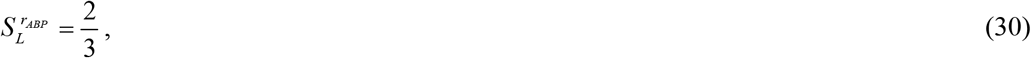

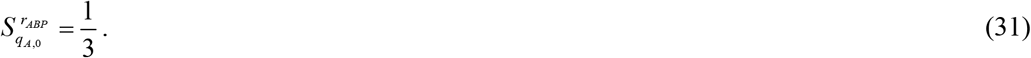

The positive values of 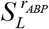 and 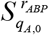 indicate that an increase in *L* and *q*_*A*,0_ will result in an increase in *r*_*ABP*_ .

Because Eq. (25) is only applicable for the infinite half-life of Aβ monomers and aggregates, the sensitivities of the growing Aβ plaque radius, *r*_*ABP*_, to *T*_1/ 2, *A*_ and *T*_1/ 2,*B*_ were computed numerically, for example:

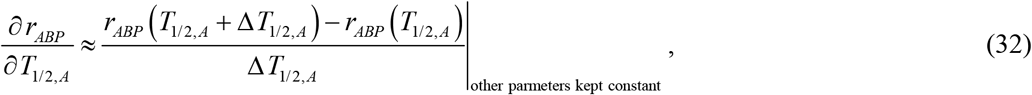

where Δ *T*_1/ 2, *A*_ = 10^−3^ *T* _1/ 2, *A*_ represents the step size. Computations were conducted with various step sizes to assess whether the sensitivity coefficients remained unaffected by changes in the step size.

## 3. Results

Figs. 2-7 were generated using the parameter values listed in Table 2, unless otherwise specified. Dysfunctional autophagic/lysosomal pathways [50,51] and impaired ubiquitin-proteasome systems [52] may potentially contribute to AD, as these defects can lead to the accumulation of misfolded proteins. For this reason, one of the scenarios investigated in the figures involves dysfunctional protein degradation machinery, which could result in infinitely large half-lives of Aβ monomers and/or aggregates.

**Fig. 2.**
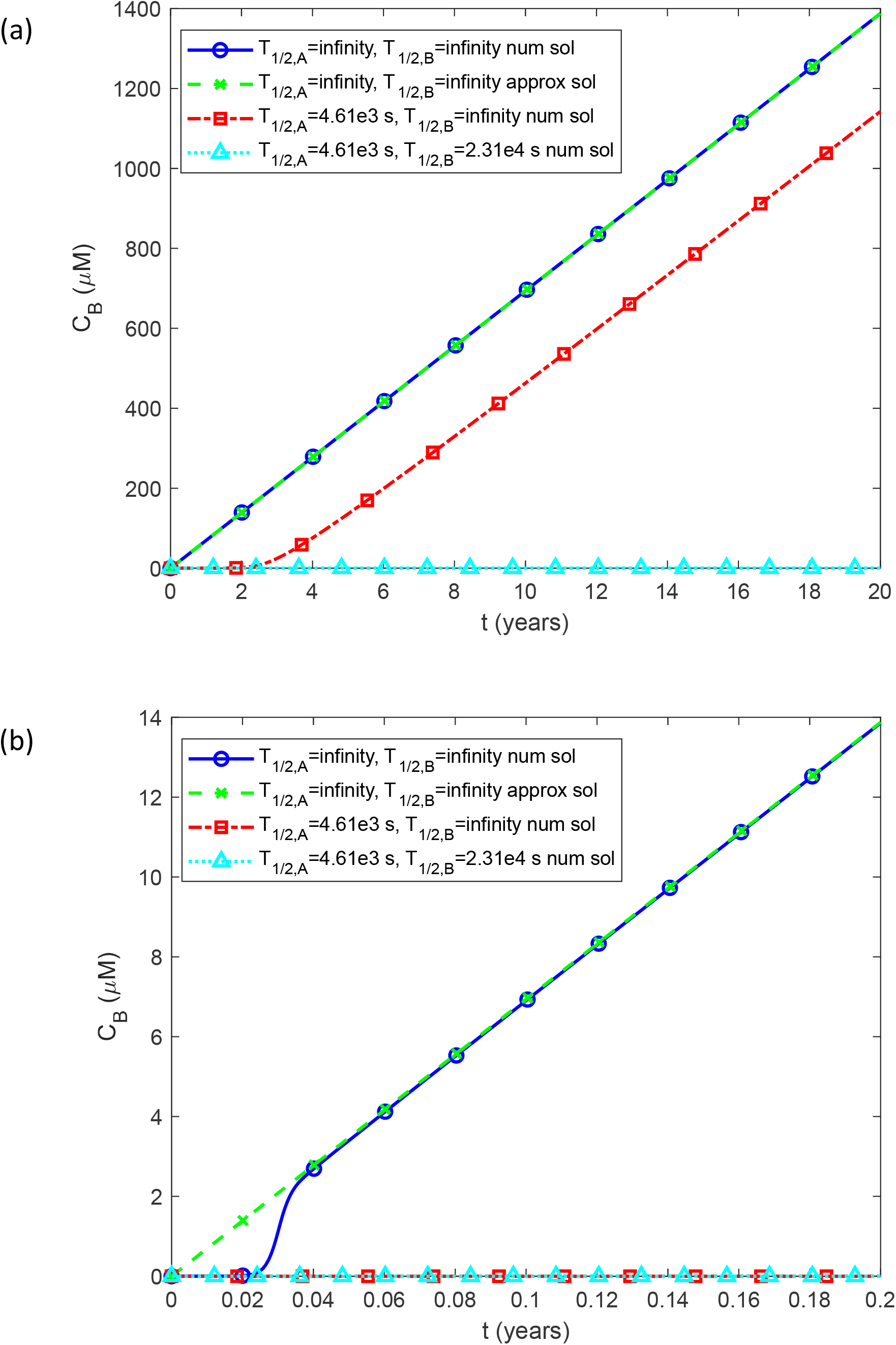
(a) Molar concentration of Aβ aggregates, *C*_*B*_, vs time (in years). (b) Similar to Fig. 2a but focusing on a specific time range of [0 0.2 year] on the *x*-axis. The numerical solution is obtained by solving Eqs. (3) and (4) numerically with the initial condition (5). The approximate solution is plotted using Eq. (17) and converting the dimensionless variables into dimensional ones. The approximate solution is applicable only for *T*_1/ 2, *A*_ ⟶∞ and *T*_1/ 2,*B*_ ⟶∞, and its plot is exclusively presented for this specific scenario. *L*=50 μm and *q*_*A*,0_ =1.1×10_^−4^_ μM μm s^-1^.

In the scenario where Aβ monomers have an infinite half-life, the concentration of Aβ aggregates, *C*_*B*_, increases linearly with time, consistent with the approximate solution provided by Eq. (17) (Fig. 2a). However, it is worth noting that this approximate solution is not valid at small times within the range of [0 0.04 year] (Fig. 2b). When the machinery responsible for degrading Aβ monomers works properly ( *T*_1/ 2, *A*_ = 4.61×10^3^ s, *T*_1/ 2,*B*_ ⟶∞ ), the computational results show an approximately 2-year delay before the linear increase in Aβ aggregate concentration begins (Fig. 2a). This suggests that functional proteolytic machinery, which degrades Aβ monomers, can postpone the formation of Aβ plaques by several years [53]. If Aβ aggregates also degrade, in addition to monomer degradation ( *T*_1/ 2, *A*_ = 4.61×10^3^ s, *T*_1/ 2,*B*_ = 2.31×10^4^ s), no aggregate growth is observed, as depicted in Fig. 2.

It should be noted that the concentrations of Aβ aggregates, denoted as *C*_*B*_, plotted in Fig. 2, are computed without considering the removal of aggregates from the cytosol because of their deposition into the amyloid plaques. The model assumes that all Aβ aggregates eventually become part of the plaques, see Eqs. (22) and (25).

In the scenario where Aβ monomers have an infinite half-life, the concentration of Aβ monomers, *C*_*A*_, decreases as time progresses (Fig. 3a,b). This decline is a consequence of the increasing concentration of Aβ aggregates (Fig. 2a), leading to faster autocatalytic conversion of monomers into aggregates as described by Eq. (4). The approximate solution given by Eq. (16) closely matches the numerical solution (Fig. 3a), but only for times greater than 0.04 years (Fig. 3b). In the scenario where the machinery responsible for degrading Aβ monomers functions normally ( *T*_1/ 2, *A*_ = 4.61×10^3^ s, *T*_1/ 2,*B*_ ⟶∞ ), there is approximately a 2-year delay in the decrease of Aβ monomer concentration. This delay is due to the postponed production of Aβ aggregates (as shown in Fig. 2a). These aggregates play a role in the autocatalytic conversion of monomers into aggregates (refer to Eq. (3)), which accelerates the monomer decay. If, in addition to monomer degradation, Aβ aggregates also degrade ( *T*_1/ 2, *A*_ = 4.61×10^3^ s, *T*_1/ 2,*B*_ = 2.31×10^4^ s), the concentration of monomers stays at a constant value independent of time (Fig. 2).

**Fig. 3.**
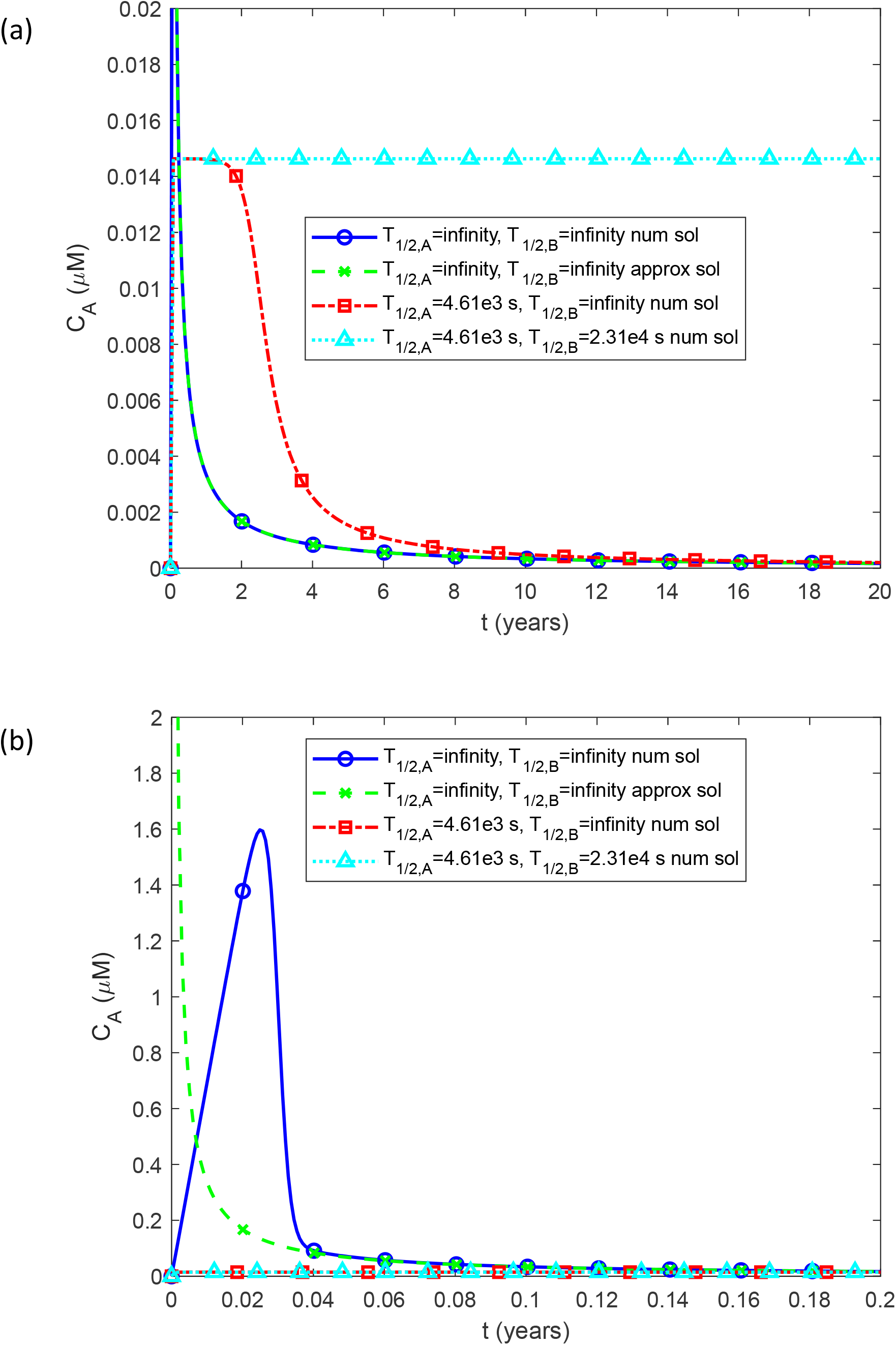
(a) Molar concentration of Aβ monomers, *C*_*A*_, vs time (in years). (b) Similar to Fig. 3a but focusing on a specific time range of [0 0.2 year] on the *x*-axis. The numerical solution is obtained by numerically solving Eqs. (3) and (4) with the initial condition (5). The approximate solution is plotted using Eqs. (16) and converting the dimensionless variables into dimensional ones. The approximate solution is applicable only for *T*_1/ 2, *A*_ ⟶∞ and *T*_1/ 2,*B*_ ⟶∞, and its plot is exclusively presented for this specific scenario. *L*=50 μm and *q*_*A*,0_ =1.1×10_^−4^_ μM μm s^-1^.

The increase in monomer concentration due to the decay of aggregates is explained by the fact that aggregates catalyze the conversion of monomers into aggregates.

In the scenario of infinite half-life for Aβ monomers, the radius of a growing Aβ plaque, *r*_*ABP*_, increases in proportion to the cube root of time, *t*^1/3^, in agreement with the approximate solution provided in Eq. (25) (Fig. 4a). The agreement between the approximate and numerical solutions is excellent, except for when *t* is within the smaller time range of [0 0.04 year] (Fig. 4b). This implies that during the early stages, the growth of the Aβ plaque is determined by the kinetic conversion of Aβ monomers into aggregates, resulting in a sigmoidal increase in the plaque’s radius, consistent with observations in [44]. Conversely, during later stages, the growth is determined by Aβ production, as assumed in the approximate solution.

**Fig. 4.**
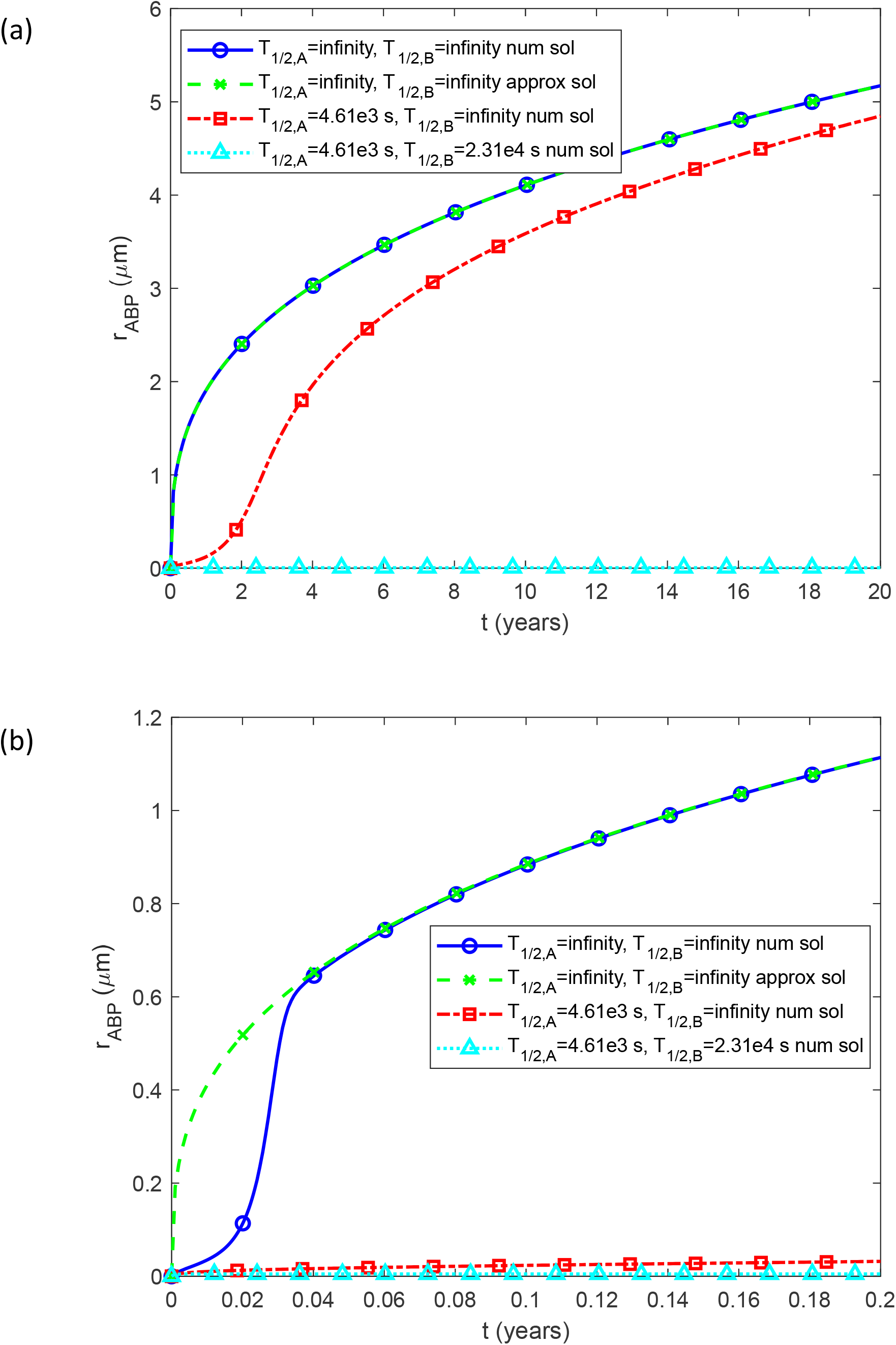
(a) Radius of a growing Aβ plaque, *r*_*ABP*_, vs time (in years). (b) Similar to Fig. 4a but focusing on a specific time range of [0 0.2 year] on the *x*-axis. The numerical solution is obtained by numerically solving Eqs. (3) and (4) with the initial condition (5) and then using Eq. (22) to calculate the radius of the Aβ plaque. The dimensionless variables are then converted into dimensional ones. The approximate solution is applicable only for *T*_1/ 2, *A*_ ⟶∞ and *T*_1/ 2,*B*_ ⟶∞, and its plot is exclusively presented for this specific scenario. *L*=50 μm and *q*_*A*,0_ =1.1×10_^−4^_ μM μm s^-1^.

If the machinery responsible for Aβ monomer degradation functions normally ( *T*_1/ 2, *A*_ = 4.61×10^3^ s, *T*_1/ 2,*B*_ ⟶∞ ), the computational result predicts a slower increase rate for the radius of an Aβ plaque over time (Fig. 4a). This suggests that the cube root hypothesis is applicable only when at least 0.04 years have passed since the beginning of Aβ aggregation, and the Aβ monomer degradation mechanism is not functioning correctly. If Aβ aggregates degrade as well ( *T*_1/ 2, *A*_ = 4.61×10^3^ s, *T*_1/ 2,*B*_ = 2.31×10^4^ s), the growth of Aβ plaque does not occur (Fig. 4).

In Fig. S1a in the Supplemental Materials, I explored how the growth of the plaque radius is affected by slower Aβ monomer degradation than physiological. All lines are computed for *T*_1/ 2,*B*_ ⟶∞ . In addition to the physiologically relevant half-life of monomers ( *T*_1/ 2, *A*_ = 4.61×10^3^ s), I studied situations where the half-life is 2x and 10x longer than physiological. I also plotted the situation corresponding to an infinite half-life of monomers ( *T*_1/ 2, *A*_ ⟶∞ ). The increase of *T*_1/ 2, *A*_ by a factor of two from physiological ( *T* _1/ 2, *A*_= 2 × 4.61×10^3^ s) brings the plaque radius halfway closer to the scenario characterized by *T*_1/ 2, *A*_ ⟶∞ . When *T*_1/ 2, *A*_ is increased by a factor of ten from physiological ( *T*_1/ 2, *A*_ = 10 × 4.61×10^3^ s), the growth of the plaque radius is almost identical to the situation with an infinite half-life of monomers (Fig. S1a).

Fig. S1b investigates how the growth of the plaque radius is influenced by slower Aβ aggregate degradation than physiological. All lines are calculated for *T*_1/ 2, *A*_ ⟶∞ s. In addition to the physiologically relevant half-life of aggregates ( *T*_1/ 2,*B*_ = 2.31×10^4^ s), I examined situations where the half-life is one hundred and ten thousand times longer than physiological. I also depicted the scenario corresponding to an infinite half-life of aggregates ( *T*_1/ 2,*B*_ ⟶∞ ). For the assumed physiological value of aggregate half-life ( *T*_1/ 2,*B*_ = 2.31×10^4^ s), the plaque radius rapidly increases to approximately 0.1 μm and then remains constant. If *T*_1/ 2,*B*_ is increased by a factor of one hundred from physiological ( *T*_1/ 2,*B*_ = 10^2^ × 2.31×10^4^ s), the plaque radius rapidly increases to approximately 0.9 μm and then remains constant. When *T*_1/ 2,*B*_ is increased by a factor of ten thousand from physiological ( *T*_1/ 2,*B*_ = 10^4^ × 2.31×10^4^ s), the plaque radius increases similarly to the situation with an infinite half-life of aggregates but remains approximately 20% below the curve corresponding to *T*_1/ 2,*B*_ ⟶∞ (Fig. S1b).

In Fig. 5, I explored a hypothetical scenario where a therapeutic intervention halts the production of Aβ monomers after 10 years. If no degradation of Aβ aggregates is assumed, plaque growth ceases after 10 years due to the absence of new monomer production (Fig. 5a). However, if Aβ aggregates are degraded (e.g., by autophagy [54]), the cessation of monomer supply results in the reduction ( *T*_1/ 2,*B*_ = 10^4^ × 2.31×10^4^ s) or complete disappearance ( *T*_1/ 2,*B*_ = 2.31×10^4^ s and *T*_1/ 2,*B*_ = 10^2^ × 2.31×10^4^ s) of the Aβ plaques (Fig. 5b).

**Fig. 5.**
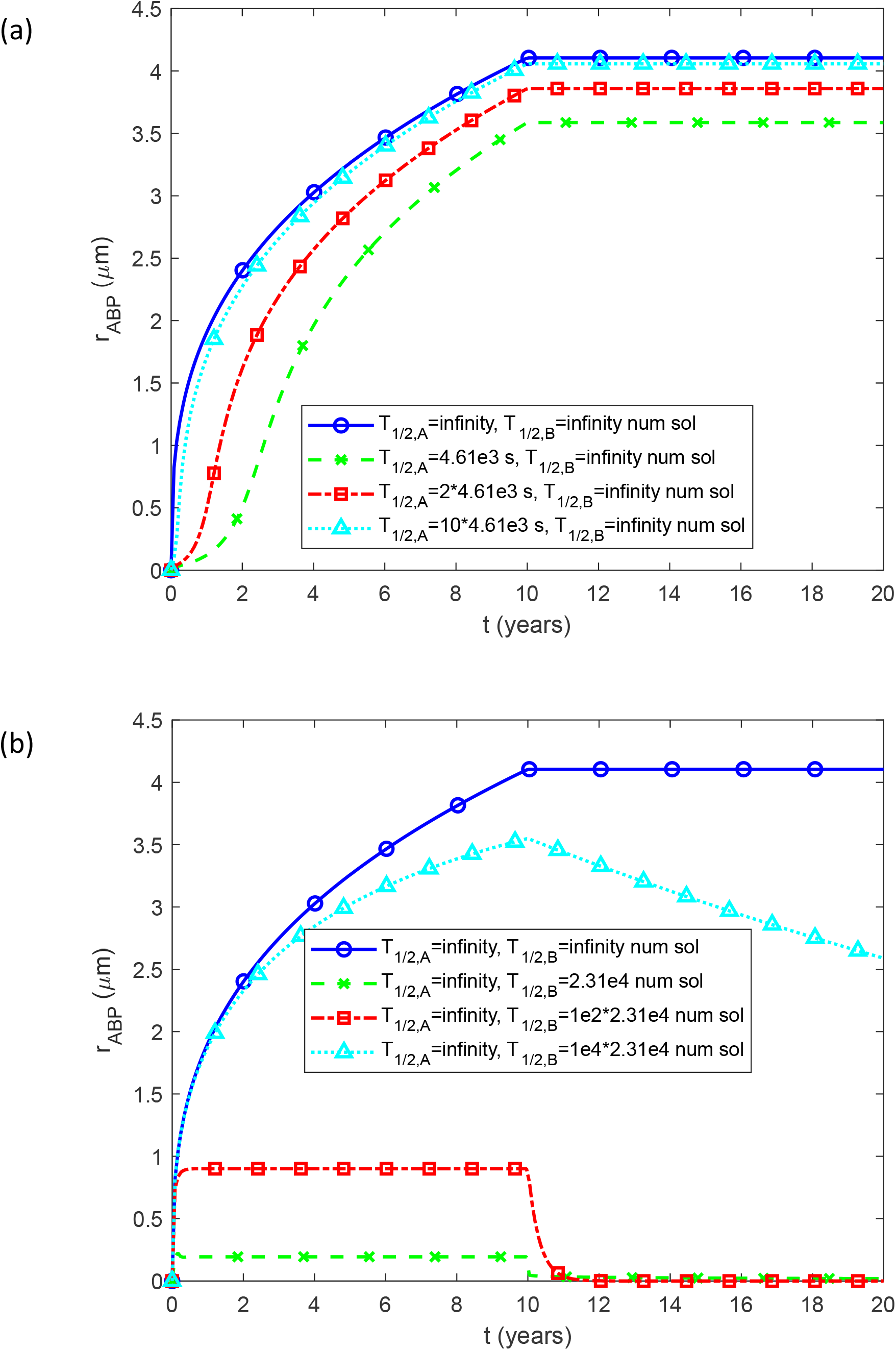
Radius of a growing Aβ plaque, *r*_*ABP*_, vs time (in years) for (a) various half-lives of Aβ monomers; (b) various half-lives of Aβ aggregates. A hypothetical situation when a therapeutic intervention causes the production of Aβ monomers to stop after 10 years. *L*=50 μm and *q*_*A*,0_ =1.1×10_^−4^_ μM μm s^-1^.

Fig. 6a illustrates the relationship between the radius of Aβ plaques after 20 years of growth and the half-distance between these plaques, *L*. When *T*_1/ 2, *A*_ ⟶∞, *r*_*ABP*_ increases with *L* almost linearly. This occurs because, for a uniform volumetric production rate of Aβ monomers, a smaller number of plaques (larger distances between them) allows each individual plaque to access a greater supply of Aβ, thus enabling it to grow to a larger size. If *L* = 500 μm, the plaque’s diameter after 20 years of growth is approximately 50 μm, which agrees with the representative diameter of an Aβ plaque observed in [32]. In the case of infinite half-life for Aβ monomers, the approximate solution provided by Eq. (25) precisely matches the numerical solution (Fig. 6a). In the scenario where Aβ monomer degradation functions normally ( *T*_1/ 2, *A*_ = 4.61×10^3^ s, *T*_1/ 2,*B*_ ⟶∞ ), an increase in *L* first results in an increase in *r*_*ABP*_, and subsequently (after reaching approximately *L* = 250 μm), it leads to a decrease in *r*_*ABP*_ (as shown in Fig. 6a). This occurs because an increase in *L* leads to a proportional increase in the supply of Aβ monomers into the CV through one of its faces, scaling with *L*^2^. The number of Aβ monomers degraded within the CV scales with *L*^3^, signifying that Aβ monomer degradation becomes the prevailing process within the CV at a certain point. If Aβ aggregates also degrade (*T*_1/2,*A*_ =4.61×10^3^ s, *T*_1/ 2,*B*_ = 2.31×10^4^ s), there is no growth of Aβ plaque, regardless of *L* (Fig. 6a).

**Fig. 6.**
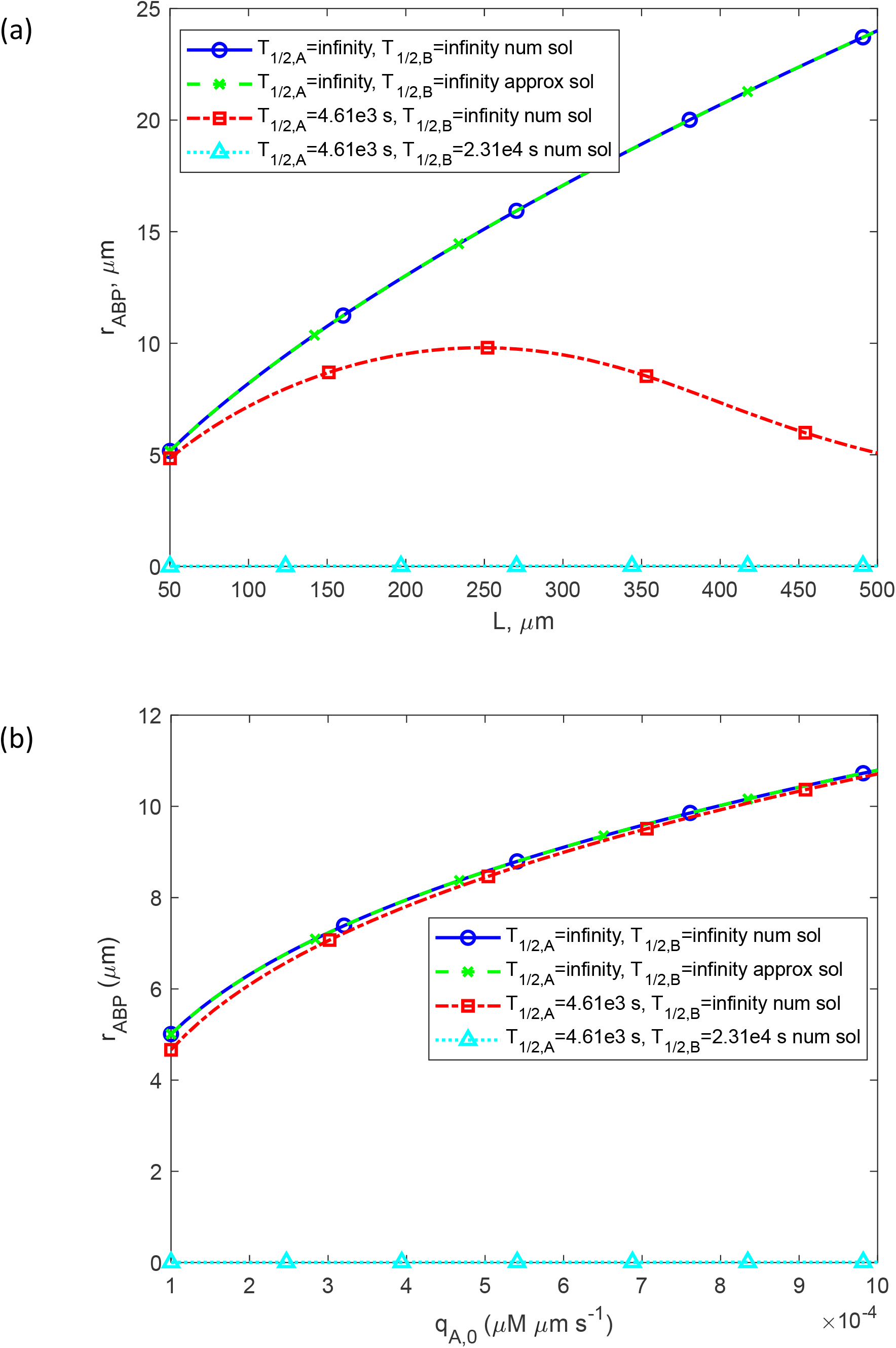
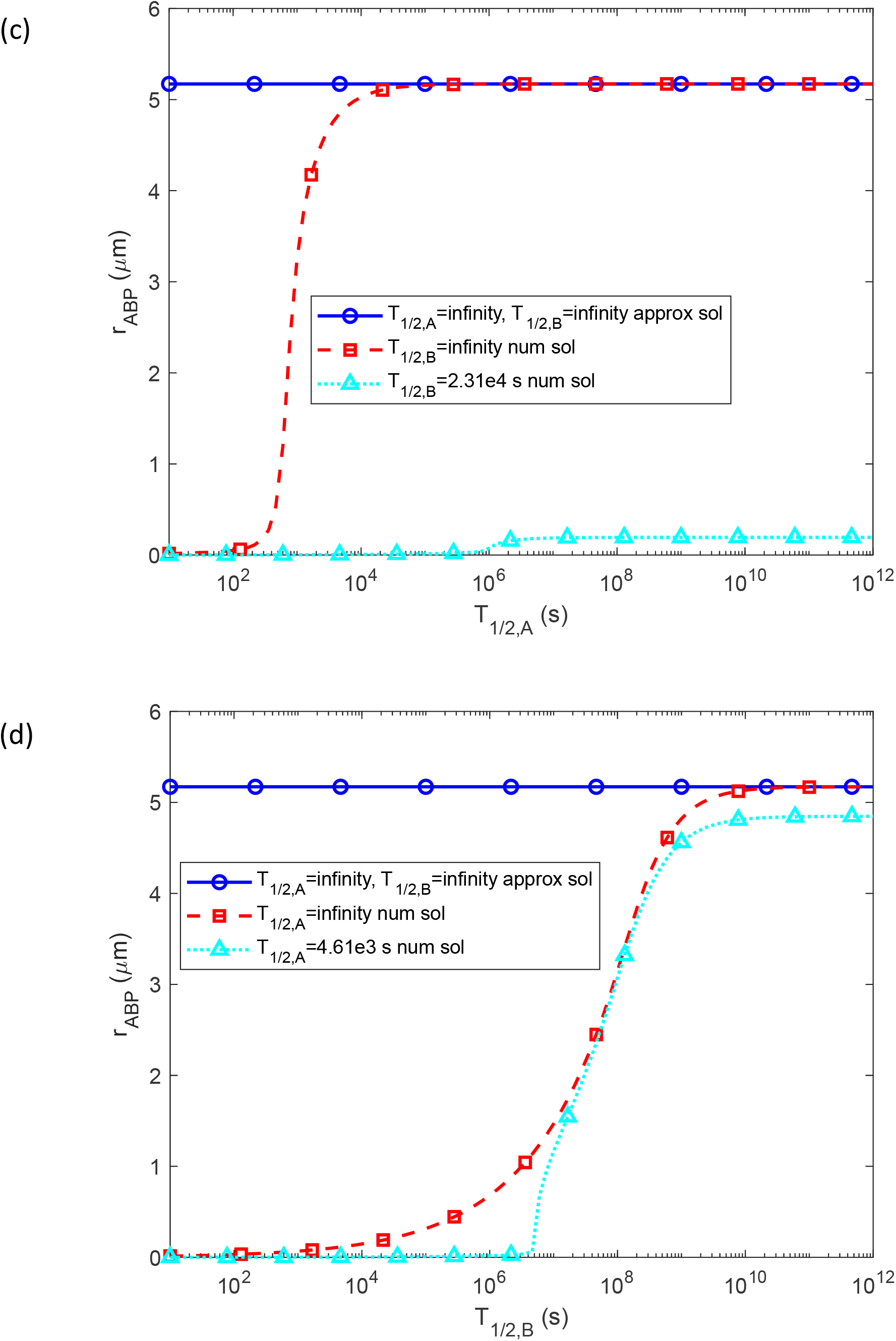
Radius of a growing Aβ plaque after 20 years of growth vs the following parameters: (a) half distance between senile plaques, *q*_*A*,0_ =1.1×10_^−4^_ μM μm s^-1^; (b) rate of production of Aβ monomers, *L*=50 μm; (c) half-life of Aβ monomers, *L*=50 μm and *q*_*A*,0_ =1.1×10_^−4^_ μM μm s^-1^; (d) half-life of Aβ aggregates, *L*=50 μm and *q*_*A*,0_ =1.1×10_^−4^_ μM μm s^-1^.

Fig. 6b illustrates how the radius of Aβ plaques after 20 years of growth is influenced by the rate of Aβ monomer production. When *T*_1/ 2, *A*_ ⟶∞ and *T*_1/ 2,*B*_ ⟶∞, *r* _*ABP*_ is directly proportional to 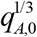 because a higher production rate of Aβ monomers results in a greater supply of Aβ available for plaque growth (Fig. 6b). In the scenario where Aβ monomers have an infinite half-life, the approximate solution given by Eq. (25) precisely agrees with the numerical solution (Fig. 6b). However, if the machinery responsible for Aβ monomer degradation is functioning normally ( *T*_1/ 2, *A*_ = 4.61×10^3^ s, *T*_1/ 2,*B*_ ⟶∞ ), the plaque grows nearly at the same rate as it does for *T*_1/ 2, *A*_ ⟶∞ . This is because Aβ monomer production is continuous, and the monomers convert into Aβ plaques before they can be broken down. If Aβ aggregates also undergo degradation (*T*_1/2,*A*_ =4.61×10^3^ s, *T*_1/ 2,*B*_ = 2.31×10^4^ s), there is no growth of the Aβ plaque, irrespective of *q*_*A*,0_ (Fig. 6b).

Fig. 6c depicts the correlation between the Aβ plaque radius after 20 years of growth and the half-life of Aβ monomers. If *T*_1/ 2,*B*_ ⟶∞ (the line marked by squares), the numerical solution produced a sigmoidal curve. For the smallest value of *T*_1/ 2, *A*_ in Fig. 6c (10 s), the plaque radius is zero. It gradually increases as the half-life of monomers increases, reaching a value corresponding to the infinite half-life of monomers at *T*_1/ 2, *B*_ = 10^5^ s. Beyond this point, it aligns with the plaque radius predicted by the approximate solution given by Eq. (25). If *T*_1/ 2, *A*_ = 2.31×10^4^ s (the line marked by triangles), the plaque radius shows no growth if *T*_1/ 2, *A*_ < 10^6^ s and very slow growth for larger values of *T*_1/ 2, *A*_, approaching approximately 1 μm for *T* = 10^12^ s (Fig. 6c).

Fig. 6d depicts the correlation between the Aβ plaque radius after 20 years of growth and the half-life of Aβ aggregates. If *T*_1/ 2, *A*_ ⟶∞ (the line marked by squares), for *T* _1/ 2, *B*_< 10^2^ s, the plaque radius is zero. It gradually increases as the half-life of aggregates increases, reaching a value corresponding to the infinite half-life of monomers at *T*_1/ 2,*B*_ = 10^10^ s. Beyond this point, it aligns with the plaque radius predicted by the approximate solution given by Eq. (25). If *T*_1/ 2, *A*_ = 4.61×10_3_ s (the line marked by triangles), the plaque radius shows no growth if *T*_1/ 2,*B*_ < 10^6^ s and S-shaped growth for larger values of *T*_1/ 2,*B*_, approaching approximately 4.7 μm for *T*_1/ 2,*B*_ = 10^12^ s (Fig. 6d).

Fig. 7 illustrates the dimensionless sensitivities of the Aβ plaque radius after 20 years of growth to various model parameters. Positive sensitivity indicates that an increase in the parameter leads to an increase in the plaque radius, while negative sensitivity signifies that an increase in the parameter results in a decrease in the plaque radius.

**Fig. 7.**
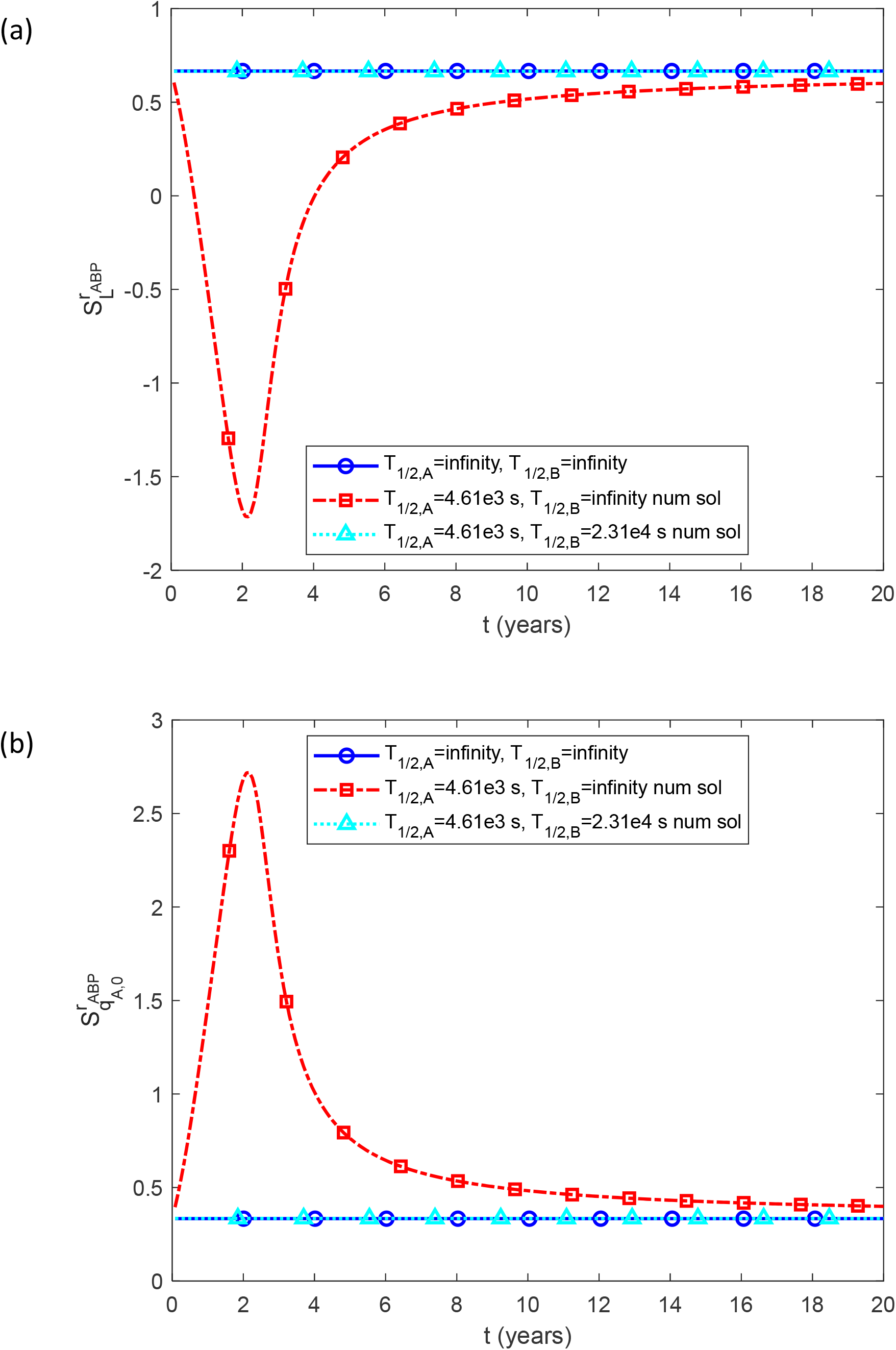
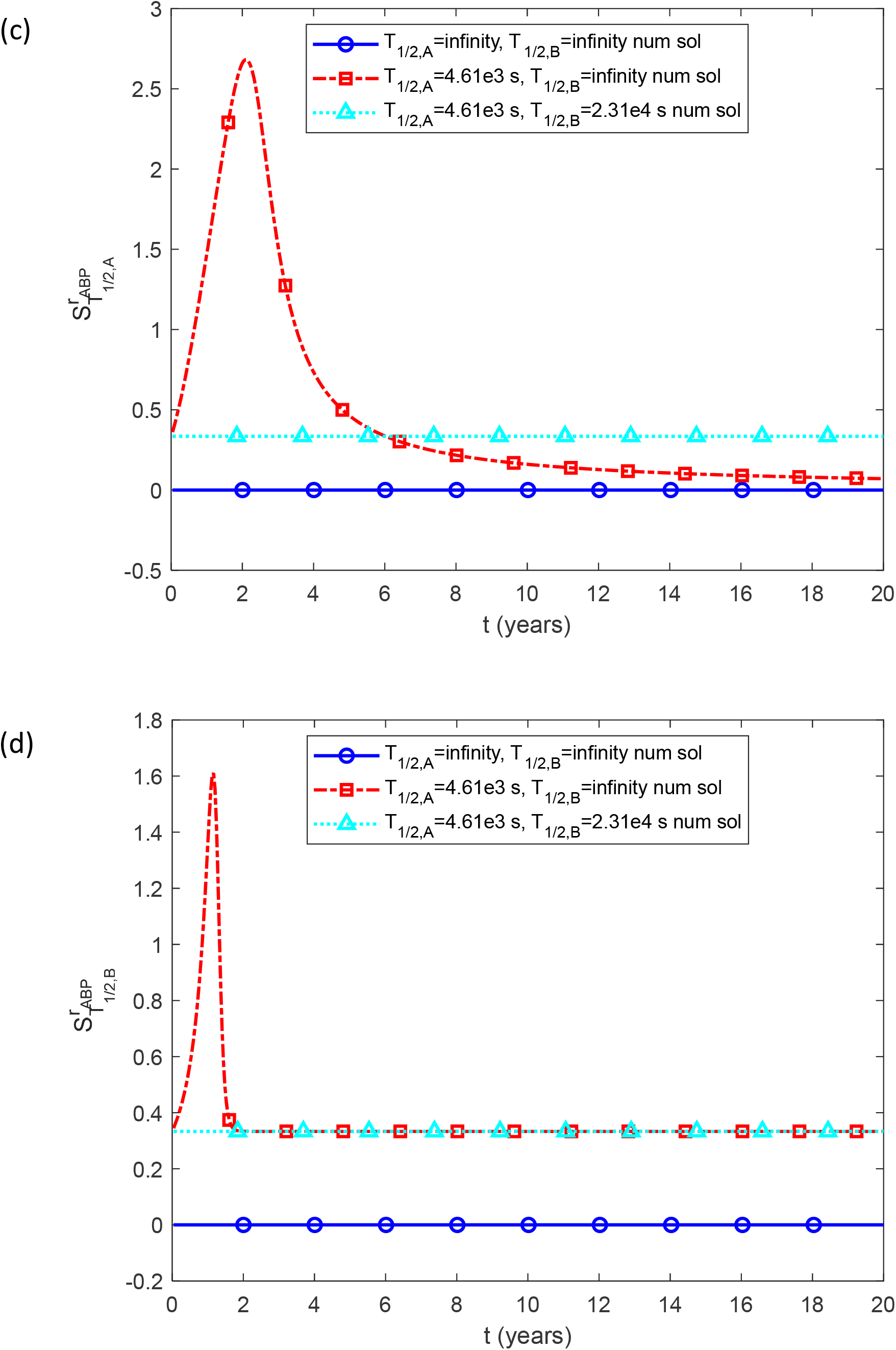
Dimensionless sensitivity of the Aβ plaque’s radius, *r*_*ABP*_, to the following parameters: (a) half distance between senile plaques, *L, q*_*A*,0_ =1.1×10_^−4^_ μM μm s^-1^, Δ*L* = 10^−3^ *L* ; (b) rate of production of Aβ monomers, *q*_*A*,0_, *L*=50 μm, Δ*q*_*A*,0_ = 10^−3^ *q*_*A*,0_ ; (c) half-life of Aβ monomers, *T*_1/ 2, *A*_, *q*_*A*,0_ =1.1×10^−4^ μM μm s^-1^, *L*=50 μm, Δ *T*_1/ 2, *A*_ = 10^−3^*T*_1/ 2, *A*_ ; (d) half-life of Aβ aggregates, *T*_1/ 2, *B*_, *q*_*A*,0_ =1.1×10^−4^ μM μm s^-1^, *L*=50 μm, Δ *T*_1/ 2, *B*_ = 10^−3^ *T*_1/ 2,*B*_ .

When the Aβ monomer degradation machinery is functioning normally ( *T*_1/ 2, *A*_ = 4.61×10^3^ s, *T*_1/ 2,*B*_ ⟶∞ ), the dimensionless sensitivity of the Aβ plaque’s radius to *L* decreases over time, turning negative after 0.5 years of growth. It reaches its minimum value after approximately 2 years of growth, then becomes positive again after 4 years of growth (Fig. 7a). In the scenario of infinite monomer half-life, the sensitivity equals 2/3, consistent with Eq. (30). If Aβ aggregates also undergo degradation (*T*_1/2,*A*_ =4.61×10^3^ s, *T*_1/ 2,*B*_ = 2.31×10^4^ s), the value of 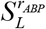 remains equal to 2/3 at any time *t* (see Fig. 7a).

The dimensionless sensitivity of the Aβ plaque’s radius to *q*_*A*,0_ is positive for *T*_1/ 2, *A*_ = 4.61×10^3^ s and *T*_1/ 2,*B*_ ⟶∞ (Fig. 7b). When Aβ monomer half-life is infinitely large, the sensitivity equals 1/3, consistent with Eq. (31). If Aβ aggregates also degrade (*T*_1/2,*A*_ =4.61×10^3^ s, *T*_1/ 2,*B*_ = 2.31×10^4^ s), the value of 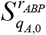 remains equal to 1/3 at any time *t* (see Fig. 7b).

At *T*_1/ 2, *A*_ = 4.61×10^3^ s and *T*_1/ 2,*B*_ ⟶∞ the dimensionless sensitivity to *T*_1/ 2, *A*_ is positive, whereas for the infinite half-lives of Aβ monomers and aggregates, the dimensionless sensitivity to *T*_1/ 2, *A*_ is zero (Fig. 7c). If both Aβ monomers and aggregates undergo degradation (*T*_1/2,*A*_ =4.61×10^3^ s, *T*_1/ 2,*B*_ = 2.31×10^4^ s), the value of 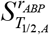 remains equal to 1/3 at any time *t* (see Fig. 7c).

At *T*_1/ 2, *A*_ = 4.61×10^3^ s and *T*_1/ 2,*B*_ ⟶∞, the dimensionless sensitivity to *T*_1/ 2,*B*_ is positive (Fig. 7d). If both Aβ monomers and aggregates are degraded (*T*_1/2,*A*_ =4.61×10^3^ s, *T*_1/ 2,*B*_ = 2.31×10^4^ s), 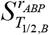 remains equal to 1/3 at any time *t* (see Fig. 7d).

Fig. 7a is computed using the step size Δ*L* = 0.001*L*, as indicated after Eq. (32). Step sizes for Fig. 7b, 7c, and 7d were selected similarly. To assess the independence of the computed sensitivity of the step size, sensitivities were computed for three different step sizes in Fig. S2. All three lines in each figure (Fig. 2a and 2b) coincide.

## 4. Discussion, model limitations, and future directions

The approximate solution predicting the radius of Aβ plaques over time suggests that these plaques increase in size following a cube root relationship with time. This result helps clarify why larger plaques exhibit slower growth. Moreover, the model indicates that the radius of Aβ plaques at large times remains unaffected by variations in the kinetic constants governing Aβ plaques aggregation rate, *k*_1_ and *k*_2_ . This observation implies that the kinetics of Aβ aggregation is not the primary limiting factor dictating Aβ plaque growth; instead, the rates of Aβ monomer production and degradation of both monomers and aggregates play more crucial roles.

The conclusion regarding the independence of kinetic rates is only applicable to the scenario with infinitely large half-lives for Aβ monomers and aggregates. It is crucial to emphasize that the approximate solution is valid only in this scenario. Notably, individuals with mutations leading to higher Aβ42 production, which has a greater tendency to aggregate than Aβ40, are more prone to AD [5,19,55].

However, these AD patients exhibit an increased aggregation-independent removal rate of Aβ42 compared to cognitively normal individuals [56], rendering the assumption of an infinite half-life for Aβ42 inapplicable.

The analysis of the exact solution of the governing equations indicates that at small times, the plaque radius depends on the kinetic constant characterizing the nucleation of Aβ aggregates. However, this dependence disappears after 0.04 years (Fig. 4b), and after that, the plaque radius grows in accordance with the approximate solution, which is independent of the kinetic constants.

Investigating a hypothetical scenario where a therapeutic intervention stops the production of Aβ monomers after 10 years reveals that if Aβ aggregates undergo degradation, the halt in monomer supply leads to either a reduction or complete disappearance of the Aβ plaques.

There are several limitations associated with this model. One limitation is the use of the lumped capacitance model to simulate the buildup of Aβ monomer and aggregate concentrations. This model assumes that these concentrations vary with time but not with location within the CV. Given that Aβ monomers are generated at lipid membranes, it is reasonable to expect concentration variations between neurons, with the highest concentration occurring at the lipid membrane. Future models should consider incorporating diffusion-driven transport of Aβ monomers between neurons, as explored in a preliminary manner in [57]. Another limitation arises from the utilization of the F-W model for Aβ aggregation, which does not distinguish between aggregates of varying sizes, such as dimers, oligomers, protofibrils, and fibrils [19]. This limitation can be addressed by employing more complex models, such as an extension of the F-W model described in [58]. This will lead to more accurate characterization of Aβ plaque structure, particularly their heterogeneity. Utilizing enhanced resolution tomography and microscopy, [32] demonstrated that Aβ assemblies comprise a dense core of higher order Aβ species, surrounded by a peripheral halo of small Aβ structures. Future models should aim at simulating this core/halo structure.

Another intriguing avenue for future research involves simulating Aβ aggregation in murine models of AD [59]. This research is important as these models do not entirely mimic human pathology, and it would aid in gaining a better understanding of the constraints of mouse models of AD.

## Abbreviations

Aβ: amyloid-β
AD: Alzheimer’s disease
APP: amyloid precursor protein
CV: control volume
F-W: Finke-Watzky

## Data accessibility

This article has no additional data. Authors’ contributions. AVK is the sole author of this paper.

## Competing interests

The author declares no competing interests.

## Funding statement

The author acknowledges the support provided by the National Science Foundation (grant CBET-2042834) and the Alexander von Humboldt Foundation through the Humboldt Research Award.

## Supplemental Materials

## S1. Supplemental figures

**Fig. S1.**
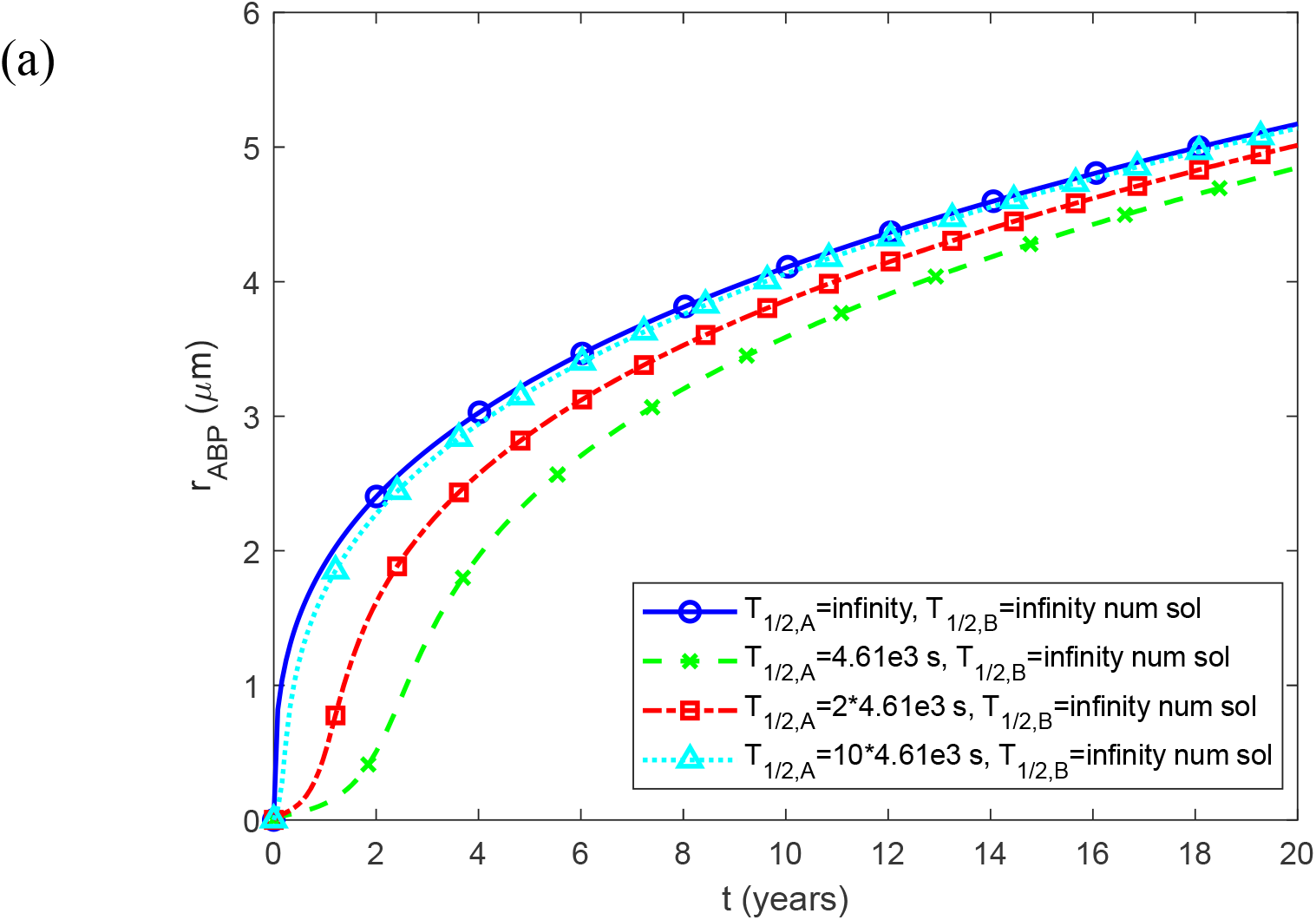
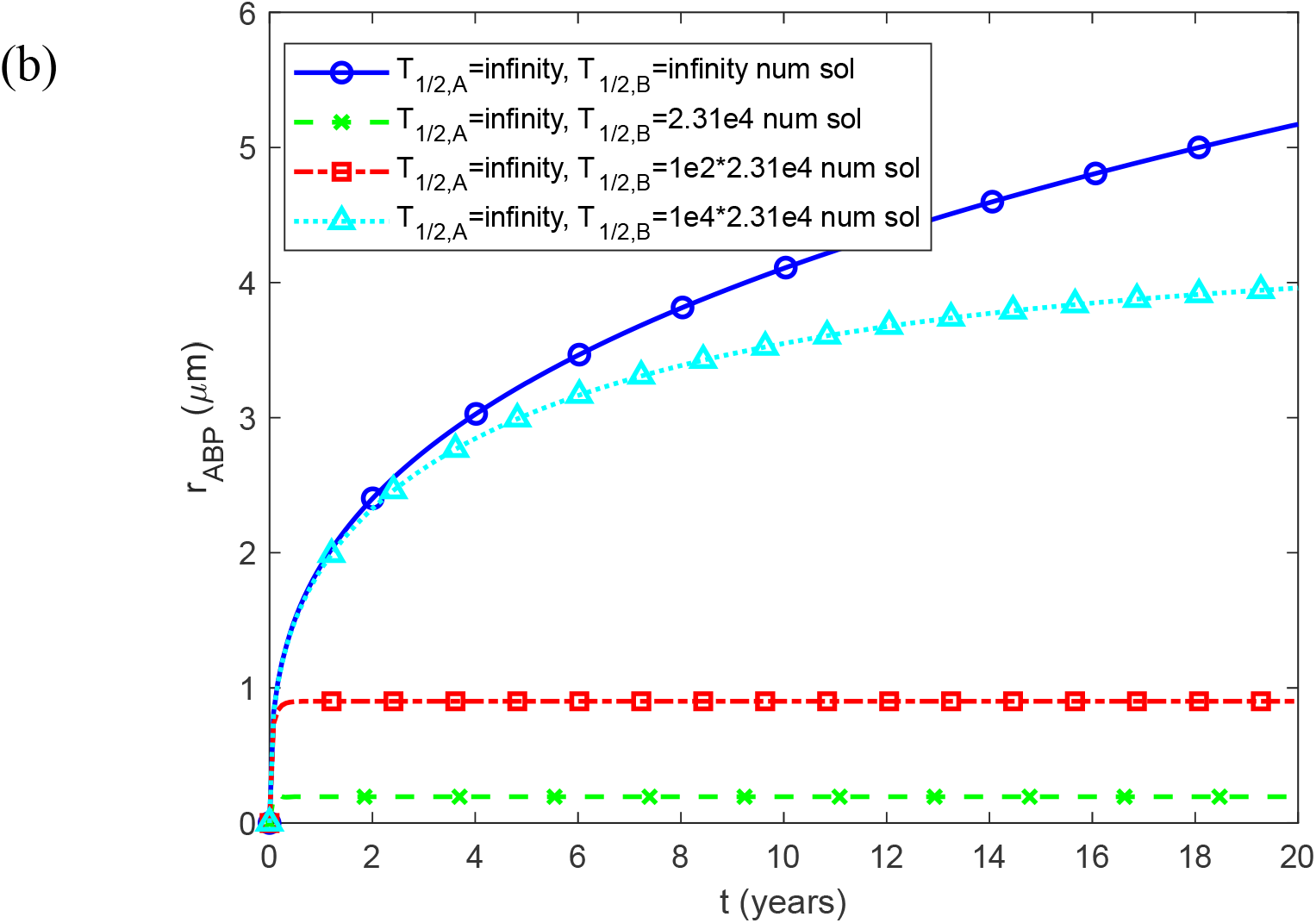
Radius of a growing Aβ plaque, *r*_*ABP*_, vs time (in years) for (a) various half-lives of Aβ monomers; (b) various half-lives of Aβ aggregates. *L*=50 μm and *q*_*A*,0_ =1.1×10_^−4^_ μM μm s^-1^.

**Fig. S2.**
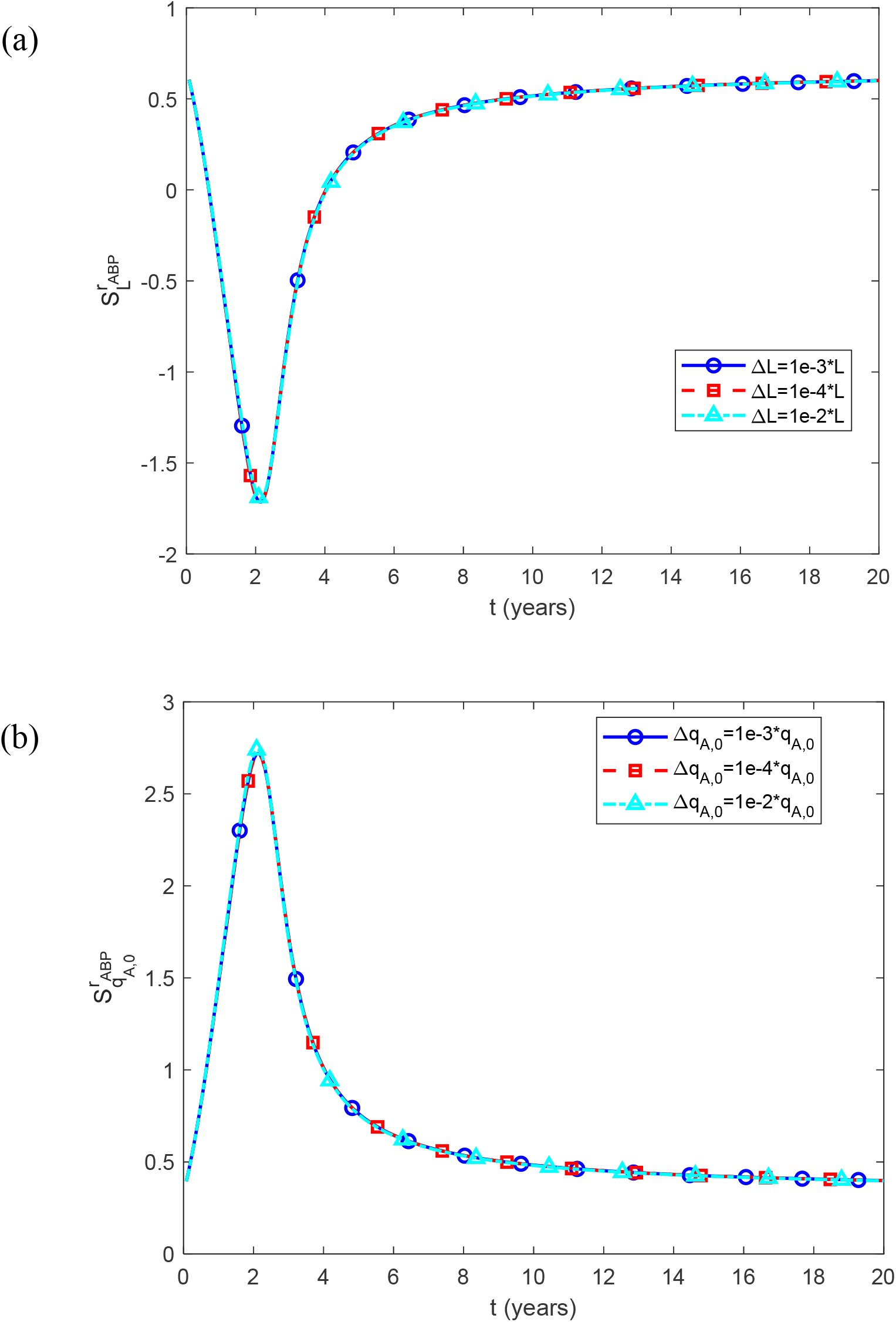
Dimensionless sensitivity of the Aβ plaque’s radius, *r*_*ABP*_, to the following parameters: (a) half distance between senile plaques, *L, q*_*A*,0_ =1.1×10_^−4^_ μM μm s^-1^; (b) rate of production of Aβ monomers, *q*_*A*,0_, *L*=50 μm. To test the independence of the computed sensitivity of the step size, three different step sizes are used in each figure. *T*_1/ 2, *A*_ = 4.61×10^−3^ s, *T*_1/ 2,*B*_ ⟶∞ .

## Notes

### Competing Interest Statement

The authors have declared no competing interest.

### Summary of Updates

Investigated a scenario where a therapeutic intervention halts the production of amyloid beta monomers after 10 years. This investigation reveals that if amyloid beta aggregates undergo degradation, the halt in monomer supply leads to either a reduction or complete disappearance of the amyloid beta plaques.

